# Stardust: improving spatial transcriptomics data analysis through space aware modularity optimization based clustering

**DOI:** 10.1101/2022.04.27.489655

**Authors:** Simone Avesani, Eva Viesi, Luca Alessandrì, Giovanni Motterle, Vincenzo Bonnici, Marco Beccuti, Raffaele Calogero, Rosalba Giugno

**Author notes:** equal contributor (Raffaele Calogero, Rosalba Giugno). equal contributor (Simone Avesani, Eva Viesi, Luca Alessandrì).

## Abstract

**Background:** Spatial transcriptomics (ST) combines stained tissue images with spatially resolved high-throughput RNA sequencing. The spatial transcriptomic analysis includes challenging tasks like clustering, where a partition among data points (spots) is defined by means of a similarity measure. Improving clustering results is a key factor as clustering affects subsequent downstream analysis. State-of-the-art approaches group data by taking into account transcriptional similarity and some by exploiting spatial information as well. However, it is not yet clear how much the spatial information combined with transcriptomics improves the clustering result.

**Results:** We propose a new clustering method, *Stardust*, that easily exploits the combination of space and transcriptomic information in the clustering procedure through a manual or fully automatic tuning of algorithm parameters. Moreover, a parameter-free version of the method is also provided where the spatial contribution depends dynamically on the expression distances distribution in the space. We evaluated the proposed methods results by analysing ST datasets available on the 10x Genomics website and comparing clustering performances with state-of-the-art approaches by measuring the spots stability in the clusters and their biological coherence. Stability is defined by the tendency of each point to remain clustered with the same neighbours when perturbations are applied.

**Conclusions:** *Stardust* is an easy-to-use methodology allowing to define how much spatial information should influence clustering on different tissues and achieving more stable results than state-of-the-art approaches.

## Background

Single-cell RNA sequencing (scRNA-seq) has emerged as an essential tool to investigate cellular heterogeneity [1]. Individual cells of the same phenotype are commonly viewed as identical functional units of a tissue or organ. However, single-cell sequencing results suggest the presence of a complex organization of heterogeneous cell states producing together system-level functionalities. Thus, the comprehension of cell transcriptomics in their morphological context is crucial to understand the effect of tissue organization in complex diseases, like specific cancer subtypes [2]. The pioneering technology called Spatial Transcriptomics (ST) [3,4,5] is able to preserve spatial information in transcriptomics, by integrating the features of microarray and the scRNA-seq barcoding system. In contrast to single-cell sequencing, spatial transcriptomics is only able to sequence the merged transcriptome profile of a small group of cells, also called a spot. By adding spatial information to scRNA-seq data, spatially resolved transcriptomes are reshaping our understanding of tissue functional organization [6]. The progressive increase in the use of ST technology highlights the need for new methods for optimizing the extraction of knowledge from ST data [7,8,9,10].

A spatial transcriptomics analysis involves upstream analysis such as data preprocessing, gene imputation and spatial decomposition, and downstream analysis such as spatial clustering, identification of spatially variable genes and genes/cells interactions. The technology is continuously evolving, raising significant challenges on the above workflow in all different steps, however, downstream analysis tends to be technology agnostic. Among all the emerging contributions in this young research area, several tools can be considered state of the art, mainly focused on downstream cluster analysis of ST data [11,12,13,14]. Pham et al. presented *stLearn* [11] to perform downstream analysis and cell types development states identification, by integrating tissue morphology, spatial dimensionality and the transcriptional information extracted from the cells. *stLearn* uses a deep neural network model to perform tile-based feature extraction from high-resolution histology images. The extracted morphological features, together with the expression value of the neighbouring spots, are exploited to smooth the gene expression data before the clustering task. Then, *stLearn* applies the Louvain or k-means clustering methods to derive the cluster to which each spot belongs. To cluster data, *stLearn* takes as input the number of principal components (PCs), the number of neighbours to build the kNN (k-nearest neighbors) graph and the resolution of the clustering algorithm.

In the same year, Hu et al. developed *SpaGCN* [12] which introduces a data integration approach based on graph convolution. *SpaGCN* as *stLearn* adds the histological information in the clustering task. It represents, through a weighted graph, the gene expression and also the similarity between each pair of spots. The latter is calculated taking into account the spatial coordinates of the spots and the average RGB value in a square of pixels to which the spots belong. The method allows increasing the weight given to histological information by varying the contribution of spots when aggregating gene expression data. To give a higher weight to images with a clear histological structure, the scaling parameter *s* can be increased when calculating the pairwise distance between spots. The hyper-parameter *l*, i.e. the characteristic length scale, can be tuned starting from the parameter *p*, which determines the percentage of total expression provided by the neighbours. The characteristic length scale determines the contribution of neighbouring spots when aggregating gene expression data by adjusting the edge weight between pairs of spots. Then, *SpaGCN* uses Louvain’s method on the aggregated output matrix from graph convolution layers to perform clustering. In addition, this method enables setting the size of the RGB square of pixels, the number of PCs and the resolution of the clustering algorithm. Moreover, users have the possibility to discard or keep the image information by setting a Boolean flag. *SpaGCN*, as other tools, allows identifying spatially variable genes (SVGs) or meta genes for each resulting spatial domain to give a biological meaning to the detected clusters as reported in [7].

Subsequently, Dries et al. [13] presented *Giotto*, a toolbox of algorithms, including a Hidden Markov Random Field (HMRF) method, to analyse spatial gene expression profiling associated with histological images. *HMRF* is a graph-based model that characterizes how many spots are influenced by the neighbours in order to assign each spot to one of *k* spatial domains, i.e. clusters, where *k* is given in input by the user. In *Giotto*, the parameters to be set are the ones given in input to the *HMRF* function, that is, the expression values to use, the name of the spatial network employed, the spatially variable genes, the spatial dimensions, the name of dimension reduction method, the number of PCs, the number of spatial domains (or clusters), three parameters (beta, tolerance and z-score) for the initialization of the method. Differently from the above methods, *Giotto* uses only spatial information of the spots and not histological information.

The same direction of *Giotto* is followed in [14] where Zhao et al. proposed a method, called *BayesSpace*, based on a Bayesian statistical approach, that improves the identification of specific profiles in tissues by imposing a Markov random field (MRF) prior that gives higher weight to spots that are spatially close. It takes as input, the number of PCs and clusters, the spatial transcriptomic platform and a series of model parameters comprising the initial cluster assignments for spots or the method to obtain the initial assignments, the error model, the precision covariance structure, the number of MCMC (Markov chain Monte Carlo) iterations, the gamma smoothing parameter, the prior mean hyperparameter, the prior precision hyperparameter and the hyperparameters for Wishart distributed precision. *BayesSpace* allows to cluster the spots according to some a priori biological knowledge or otherwise using the elbow plot of the pseudo-log-likelihood to infer the number of clusters *q* that are given in input to the method. The authors show that *BayesSpace* outperforms, in terms of adjusted rand index and manual annotations, other methods in the literature, in particular, the three widely used non-spatial algorithms, namely k-means, mclust and Louvain’s methods, and the two spatial clustering algorithms, *HMRF (Giotto)* and *stLearn*, on distinct samples of dorsolateral prefrontal cortex (DLPFC) dataset. This dataset was not analysed in our comparisons due to the lack of publicly available reference manual annotation.

In this article, we propose a downstream ST cluster method, called *Stardust*, which takes into account both the expression and the physical location in the tissue section of the transcriptional profiles, to define the similarity of the objects to be grouped. Our proposed method fits on the downstream task to perform clustering on gene count and spot location matrices. With *Stardust*, we intend to investigate how much the spatial information combined with transcriptomics improves the clustering results. *Stardust* is based on the *Seurat* [15] algorithm for the clustering of scRNA-seq data which uses Louvain’s method to perform clustering. By setting a parameter, the user can easily determine how much spatial information should affect the clustering similarity. Such a parameter can also be automatically derived from a tuning procedure. Moreover, we propose a version of *Stardust* parameter-free, *Stardust\**, i.e. it uses a dynamic non-linear formulation that changes the spatial weight according to the transcriptomics values in the surrounding space. Running time of the two methods is equivalent.

To understand how the usage of spatial information affects the stability of clusters, we evaluated both *Stardust* approaches with and without considering space on five publicly available 10x Genomics Visium datasets, respectively derived from human breast cancer section 1 (HBC1) and section 2 (HBC2), mouse kidney (MK), human heart (HH) and human lymph node (HLN) tissues [16]. We investigated the scalability of *Stardust\** testing it also on Seq-scope and Slide-seq datasets [17-18], which provide higher resolution and number of captured cells with respect to Visium datasets that combine spatial information on a tissue section with whole transcriptome sequencing at resolution of 55 µm. In Seq-scope, two sequencing steps are performed respectively allowing to retrieve spatial coordinate and captured cDNA information. We analysed two gastrointestinal tissues, liver and colon, for which the sequencing data was produced in 1 mm-wide circular areas called “tiles”, achieving a submicrometer resolution (0.5-1 μm). As for Slide-seq, we analysed the new Slide-seq V2 mouse cerebellum dataset, which consists of spatially resolved expression data at approximate resolution of single-cell (10 µm). We compared *Stardust* and *Stardust\** with currently available ST clustering methods, including *stLearn, SpaGCN, Giotto* and *BayesSpace*. Each tool comes with specific parameters to be set by the user. We fully exploited the cluster resolution parameter and used the author suggested values for all the remaining ones. In order to assess clustering performances, we exploit functional aspects such as spatially variable genes, and alternatively from current contributions, we defined two different objective clustering quality measures: the cell stability score (CSS) [19] and the coefficient of variation. The CSS defines the tendency of a cell or spot to remain clustered with the same group of elements when inducing a perturbation to the dataset, for instance by removing a random set of items, while the coefficient of variation value is derived from the CSS distribution as the ratio of the standard deviation to the mean, thus, a lower coefficient of variation means higher average stability and less variation from the mean. These comparison measures enable us to estimate the clustering stability of the different configurations and to assess whether considering spatial or morphological information leads to an improvement in terms of stability. By computing the Moran’s I [20] for each gene, we showed that genes with the highest spatial autocorrelation values, colocalize in clusters achieved by the proposed methods. Moreover, when cell type annotation is available, we verified that *Stardust* clustering maintains biological significance observing that cluster shapes appear consistent with the provided annotation.

Results show that *Stardust* and *Stardust\** improve in a statistically significant manner the clustering stability by combining the transcriptional similarity of the spots with their spatial localization in several datasets with respect to *stLearn, SpaGCN* and *Giotto*, and it is comparable with *BayesSpace*. Furthermore, while other methods force spots to form misleading cluster structures, in which neighbouring spots are clustered without sensibly sharing their expression profile, the proposed methods appear to be unaffected by such behavior. Finally, unlike other approaches, besides the number of principal components and clustering resolution parameters, *Stardust* requires only one parameter, i.e. the spatial weight, to be set by the user, and *Stardust\** does not require any parameter.

## Methods

In this Section, we introduce the proposed ST cluster approaches and the measures used for evaluating performance. Datasets used for the clustering evaluation were downloaded from the 10x Genomics website [16], respectively derived from two serial sections of human breast cancer (HBC1 and HBC2), mouse kidney (MK), human heart (HH) and human lymph node (HLN) and from the Deep Blue Data platform [21], respectively collected from colon and liver *TD*. These data together with Slide-seq V2 cerebellum dataset [18], were used to estimate method time scalability. For each 10x dataset, we loaded the associated *Seurat* object and extracted the spot coordinates and the expression matrix. In order to reduce memory usage and computation time, we filtered in the data matrices those genes expressed in more than 10 spots. To analyse Seq-scope data, we downloaded the processed RDS data files and selected a single tile for each of the two datasets chosen, specifically, the tile ID 2110 for the colon dataset and the tile ID 2117 for the liver *TD* dataset. The files containing the digital gene expression matrices and the pixel coordinates within the Slide-seq dataset were downloaded from the Single Cell Portal website referenced in Cable *et al*. [18]. Preprocessed 10x data, software code and tool documentation are available at https://github.com/InfOmics/stardust/.

### Stardust

*Stardust* is implemented on top of the *Seurat* [15] clustering algorithm. *Seurat* package is one of the most used software for scRNAseq data analysis. *Seurat* implements a network-based clustering method called the Louvain algorithm [22] which encodes each element of a dataset as a node in a graph. Pairs of nodes are connected according to a pairwise measure of similarity based on the Euclidean distance between transcriptional profiles. Then, the algorithm performs a community detection step over the graph to retrieve the dataset partition. In *Stardust*, the distance matrix used in *Seurat* is replaced with a summation of two other matrices representing the transcriptional information and the spatial position of the spots. Additionally, the distance among pairs of nodes is computed on the vectors of the distances of each node to all other nodes. The matrix regarding the transcriptional information is obtained from the pairwise Euclidean distance between transcriptional profiles in PCA space [23]. The matrix regarding the spatial position represents the pairwise spatial Euclidean distance between spots.

Given the distance matrix based on transcriptional profiles, T, and the distance matrix based on spot coordinates, S, a preliminary linear scaling step is applied to *S* in order to mitigate cases in which one measure overpowers the other. The scaling formula is the following one:

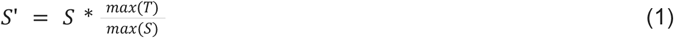

where *max(T)* and *max(S)* are the maximum value in the matrices *T* and *S*, respectively.

The user can choose between two different variants of the designed tool, namely *Stardust* or *Stardust\**, for the computation of the final distance matrix *ST. Stardust* computes the Louvain edge weights through a linear formulation and requires a fixed a priori parameter, while *Stardust\** computes the final distances through a non-linear formulation without any a priori parameter to be set.

The first method allows the user to specify a parameter called *spaceWeight*, a real number in [0,1], that defines how much to weigh the space with respect to the transcriptional similarity. By configuring a single parameter the user can control how much the space-based measure weights on the overall measure. The formula for *ST* is:

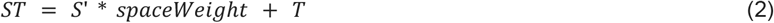

The second method first computes the normalized values of the expression distances distribution by applying the following formula:

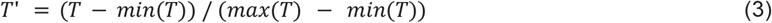

Once that *T’* is obtained, the final distance matrix *ST* is computed as a mixture of space and transcript information. The latter is always considered in its integrity, while space information is weighted by the normalized expression distances. The formula for *ST* is:

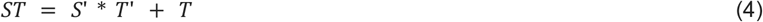

The proposed methodology is very simple and flexible, indeed it can be incorporated into any existing clustering method. Methods are developed as a standalone R package and can be easily installed from GitHub repository or used through the dedicated docker image.

### Cluster Validation

In order to give a quantitative performance evaluation of the clustering obtained, we use three different clustering quality measures: the *cell stability score* (implemented in the rCASC package) [19], the *coefficient of variation a*nd the *Moran’s I* (index) [20]. Finally, we investigate the *statistical validation* of the results.

#### Cell stability score

*rCASC* [19] takes as input a spatially resolved transcriptome and a clustering algorithm. It computes for each basic element of the dataset a *cell stability score (CSS)* that describes how much each element tends to remain clustered with the same other elements through a series of n repetitions of the clustering method on *n* different permutations of the dataset. The basic concept of the rCASC notion is that a good clustering should remain stable if a perturbation is applied to the dataset. A CSS is a real number in [0,1] associated with each individual spot in a dataset and computed running the following three steps. First, the desired clustering method is applied to the dataset and the cluster identity associated with each object - i.e. each spot in our application - are defined. Then, a subset of objects is removed from the original dataset (the percentage of objects is decided by the user) and is clustered. This step is repeated *n* times (*n* is a user parameter). We decided to set the number of permutations to 80 and to remove at each permutation 10% of the spots. In each of the repetitions, the percentage of spots that remain clustered with a particular spot in that permutation is determined by taking the results obtained in the first step as a reference. This value is stored. Finally, for each spot, it is computed how many times the set of spots clustered with it in each of the repetitions in the second step is equal to the set in the first step. This quantity is divided by the number of repetitions, obtaining the stability score. To reduce the computation time, we set a limit of 4 hours for the computation of the CSS for each tool configuration compared.

#### Coefficient of variation

To decide which configuration of *Stardust* or *Stardust\** was the best performing (based on stability scores) on a particular dataset with respect to the one not considering the space, we used the coefficient of variation defined as 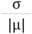, where σ is the standard deviation of the distribution of the spot scores and µ is its mean. The lower the coefficient of variation, the better the performances. We also applied this metric to the other compared methods to evaluate the best performing configuration of each tool.

#### Spatial autocorrelation of cluster markers

To evaluate the biological coherence of the obtained clusters we compute genes’ spatial autocorrelation by using the Moran’s I [20]. Moran’s I uses both feature locations and feature values simultaneously. Spatial autocorrelation is defined as a territorial cluster of similar marker values. If similar expression values of the genes are spatially localized, there is a positive spatial autocorrelation of the data. On the contrary, a spatial proximity of dissimilar values indicates a negative spatial autocorrelation. Moran’s I is defined as:

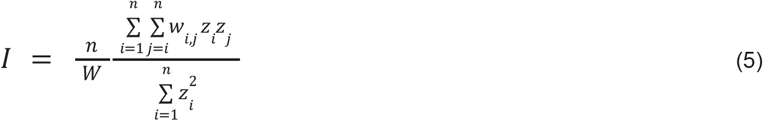

where *z*_*i*_ is the deviation of the genes from the mean, *w*_*i,j*_ is the spatial weight between observations, *n* is the number of spatial units and *w* is the sum of all *w*_*i,j*_.

#### Statistical validation

We provide statistical evidence of the variation of cluster stability achieved by the methods in the following way. Given a dataset we first apply *Stardust* or *Stardust\**, obtaining a set of stability scores i. Then, for 100 times, we shuffle the spots coordinates and re-apply the method, obtaining 100 sets of stability scores *j*_*1*_ … *j*_100_. For each couple (i,*j*_*k*_) - with k in 1…100 - we evaluate the Wilcoxon statistical test with the null hypothesis that the distribution i is greater than *j*_*k*_, i.e. the gain in stability obtained from the original spot coordinates is greater than the gain obtained from shuffled spot coordinates.

## Results

In this Section, we assess the performance of *Stardust* and *Stardust\** on five datasets from 10x Genomics, respectively derived from two serial sections of human breast cancer (HBC1 and HBC2), mouse kidney (MK), human heart (HH) and human lymph node (HLN) (see Figure 1). The dimensions of datasets are 3.798, 3.987 4.247, 4.035, 1.438 spots, respectively. Visually, the datasets show different levels of structures, from high levels such as breast tissues where we expect to find more well characterized, i.e stable, clusters to low ones as in human heart tissue.

**Figure 1:**
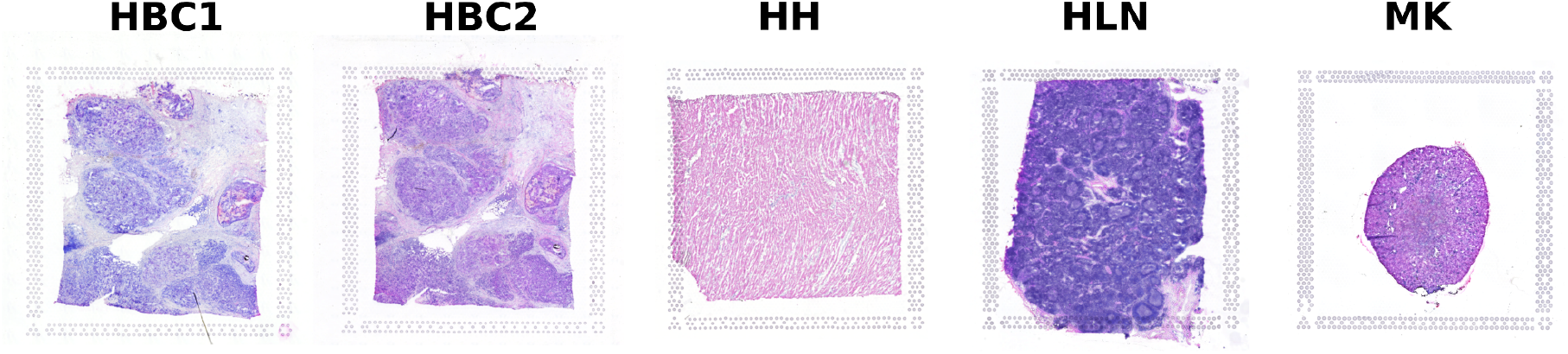
Hematoxylin and eosin (H&E) stained tissue sections of human breast cancer section 1 (HBC1), human breast cancer section 2 (HBC2), mouse kidney (MK), human heart (HH) and human lymph node (HLN).

To understand how the usage of spatial information affects the stability of clusters we computed the *cell stability scores* and evaluated the *coefficient of variation* (CV) of the stability scores (see Section Methods-Cluster Validation) of *Stardust* by varying the clustering resolution and the space weight parameters and *Stardust\** by varying only clustering resolution (see Section *Methods*). For *Stardust*, the space weights were set to 0, 0.25, 0.5, 0.75 and 1 and cluster resolution to 0.6, 0.8, and 1. Space weight equals 0, when space is not considered, corresponds to comparing *Stardust* with respect to its transcriptomic only based approach, here referred with terms *no space* used. Space weight equals 1 means that space and transcripts contribute in the same way.

Results show that the introduction of spatial information (Figures 2 (a)) reduces the coefficient of variation of each *Stardust* configuration with respect to the configuration where space is not considered. Since setting *Stardust* clustering parameters could be challenging, we also used the R package GenSA [24] solution to estimate the best combination of space weight and clustering resolution maximizing the average cell stability score. We created a dedicated Docker image where GenSA runs the *Stardust* algorithm several times to tune all the required parameters. Coefficients of variation obtained from tuned parameters are shown in Figures 2 (a) with violet dots. To limit the computation time, *Stardust* tuning was run for each dataset, fixing to 10 the maximum number of GenSA iterations. Despite the low number of iterations, the estimated average cell stability scores are all higher or comparable with our best results. The achieved CVs confirmed the trends observed from the manual tests, allowing to explore *Stardust* configurations not considered before. Tuning running time varied from 4 to 24 hours, depending on many factors, including the dataset size and computational resources. By increasing the size of the datasets or the number of combinations of parameters, GenSA does not scale and therefore it is not straightforwardly applicable on the other ST algorithms that are far more complex than *Stardust* in terms of the entire set of parameters that can be configured. A similar behavior to the one described for *Stardust* is reported for *Stardust\** in Figures 3 (a), where the dynamic setting of the spatial weight leads to a cluster resolution with an average reduced or comparable coefficient of variation to clustering without considering space.

**Figure 2:**
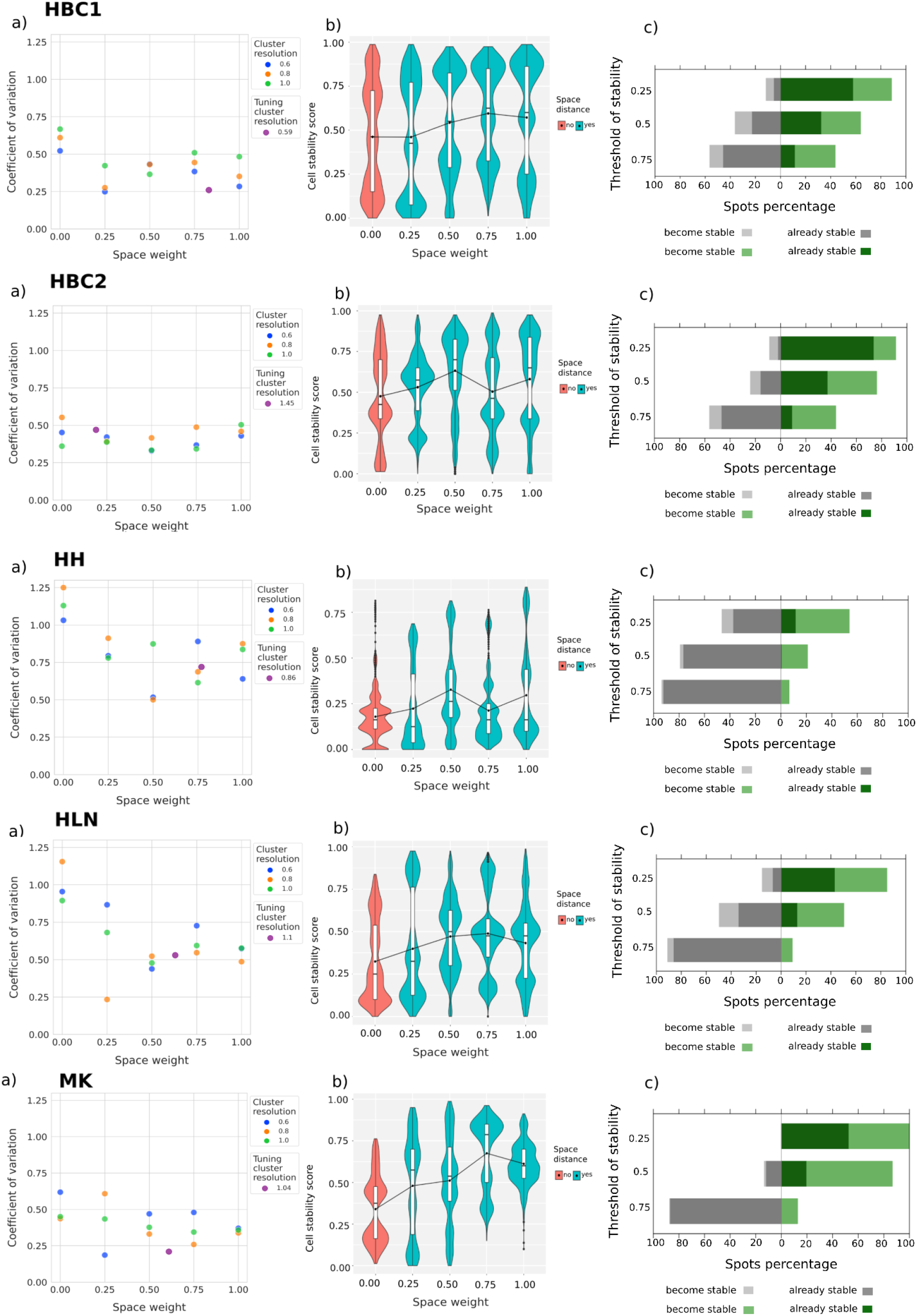
*Stardust* performance on five ST datasets: two serial sections of human breast cancer (HBC1 and HBC2), mouse kidney (MK), human heart (HH) and human lymph node (HLN). (a) *Stardust* coefficient of variation for each configuration obtained varying the space weight and clustering resolution. Space weight and resolution tuned by maximizing the average cell stability score are shown with violet dots. (b) Stability scores comparison for 5 *Stardust* configurations with increasing space weight and cluster resolution fixed to 0.8. (c) The count of spots shifting from stable to unstable and vice versa at stability thresholds equal to 0.25, 0.5, 0.75, which set the limit to consider a spot stable (above the threshold) or unstable (below the threshold), comparing the best configuration of *Stardust* (i.e., with the lowest coefficient of variation) with the one not using space information.

**Figure 3:**
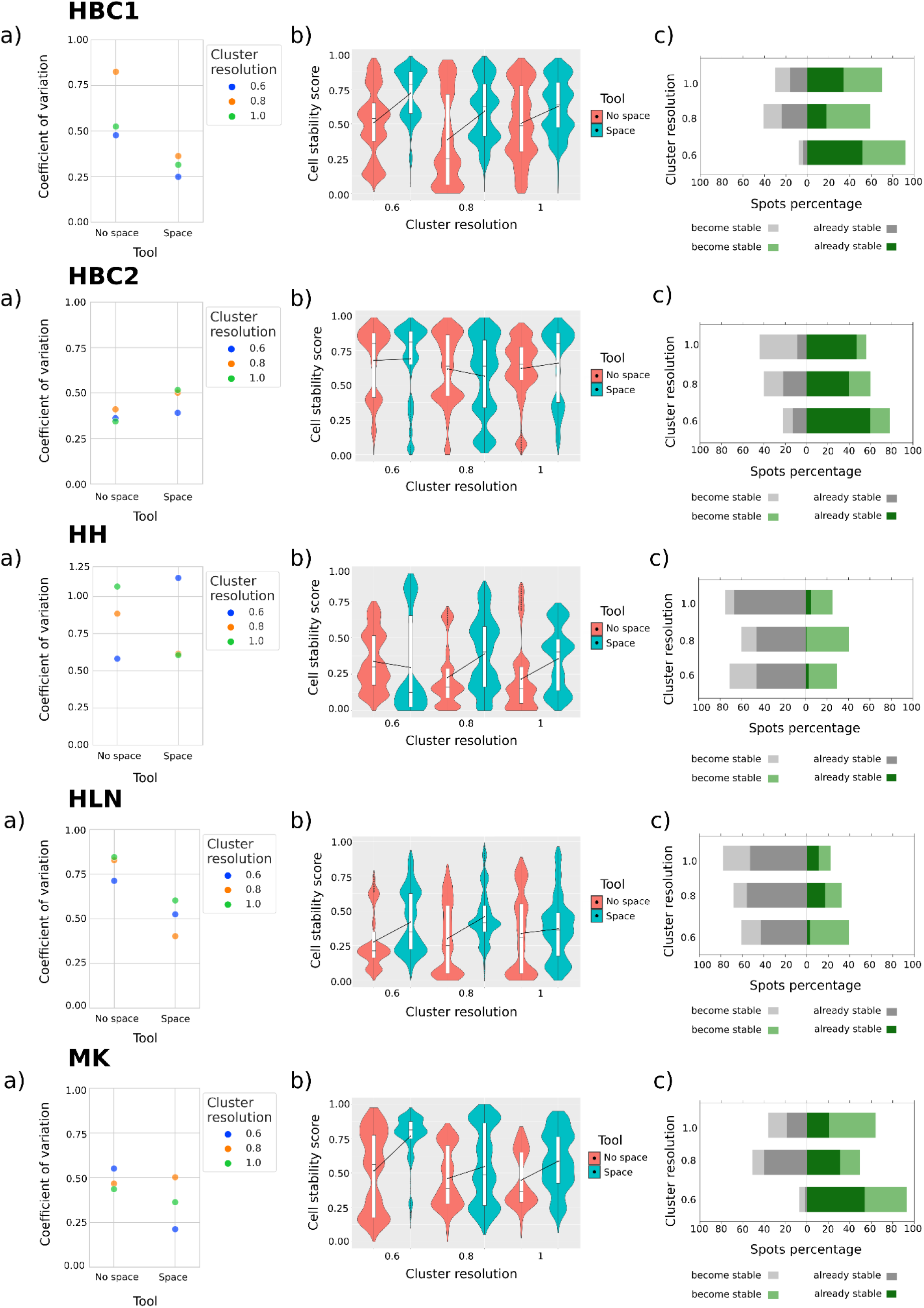
*Stardust\** space version performances with respect to *no space* version ones evaluated on five ST datasets: two serial sections of human breast cancer (HBC1 and HBC2), mouse kidney (MK), human heart (HH) and human lymph node (HLN). (a) Coefficient of variation values comparison for 3 *Stardust\** space and *no space* configurations obtained by varying the clustering resolution. (b) Stability scores comparison for 3 *Stardust* space and no space* configurations obtained by varying the clustering resolution. (c) The Stardust* count of spots shifting from stable to unstable and vice versa considering clustering with *no space* information as baseline at different clustering resolutions equal to 0.6, 0.8, 1, which set the limit to consider a spot stable (above the threshold) or unstable (below the threshold).

Figures 2-3 (b) show cluster stability improvements. Figures 2 (b) investigate five different *Stardust* space weight configurations and keep the cluster resolution fixed to 0.8, i.e. it focuses the attention on one of the *cell stability score* distributions tested in Figures 2 (a). In all the datasets, space is able to increase the overall stability, and although this behaviour is not monotonous with the increasing of the weight given to the space, different space weights allow to achieve the best scores. Figures 3 (b) show *Stardust\** cluster stability comparing its versions with and without space by varying clustering resolution. Wilcoxon test (see Section *Methods*-*Cluster Validation*) was used to evaluate the significance of the results confirming that the increase of stability scores is not due to chance.

To complete method evaluation, we compute the percentage of spots becoming stable or unstable. Figures 2 (c) compare one of the best performing configurations of *Stardust* according to the coefficient of variation values in Figures 2 (a) with the one not using space information for each dataset. Regardless of which threshold is used, the number of spots that become stable in *Stardust* with respect to the *no space* configuration is always more than the number of spots that become unstable for each dataset. However, from a cluster quality point of view, threshold values are reasonable if belonging in [0.5, 1], i.e. it is desired that each spot remains clustered with the same others in at least half of the rCASC permutations. Using the threshold 0.5, Figures 3 (c) compare *Stardust\** and the same method with *no space* information by varying the clustering resolutions, confirming that besides cluster resolution the number of spots that become stable using the space information is always more than the number of spots that become unstable for each dataset.

In Supplementary file Figures S1-5 we depict how clusters are arranged in the 2D space of the tissue section and as space information influences the clusters across the 5 *Stardust* configurations. In Figures S1-5 (a) all points are displayed, while in Figures S1-5 (b) only points with stability scores greater than or equal to 0.5 are displayed. Score values are in [0, 1], so the threshold 0.5 means that in at least half of the permutations a spot remains clustered with the same other spots and can be considered a stable one. Figures S1 and S2 show that using space the number of spots that become stable and the number of spots recognized as a unique cluster are maximized without creating structure where it’s not present as in Figure S3. In Figures S4-5 spatial information increases the overall stability of neighbouring clusters or clusters with distant spots but high transcriptional similarity. Analysing clustering results of the most stable configurations, we observed that the cluster structures reflect the biology of the tissue. In Figure 4 (a) we report a manual annotation of HBC1 from [2] and the clusters obtained from the most stable configuration of *Stardust*. Stardust\** clustering mirrors the general tissue structure allowing the identification of tumoral regions including *ductal carcinoma in situ* regions corresponding to clusters 9 and 12, and *invasive carcinoma* regions like clusters 1, 2, 4, 5, 10 and 11. Moreover we computed the Moran’s index for each HBC1 feature to identify spatial autocorrelated genes. Analysing the first 100 genes with highest Moran’s I we noticed that they colocalize in the identified clusters (see Figure 4 (b)), confirming the biological validity of *Stardust\** clustering.

**Figure 4:**
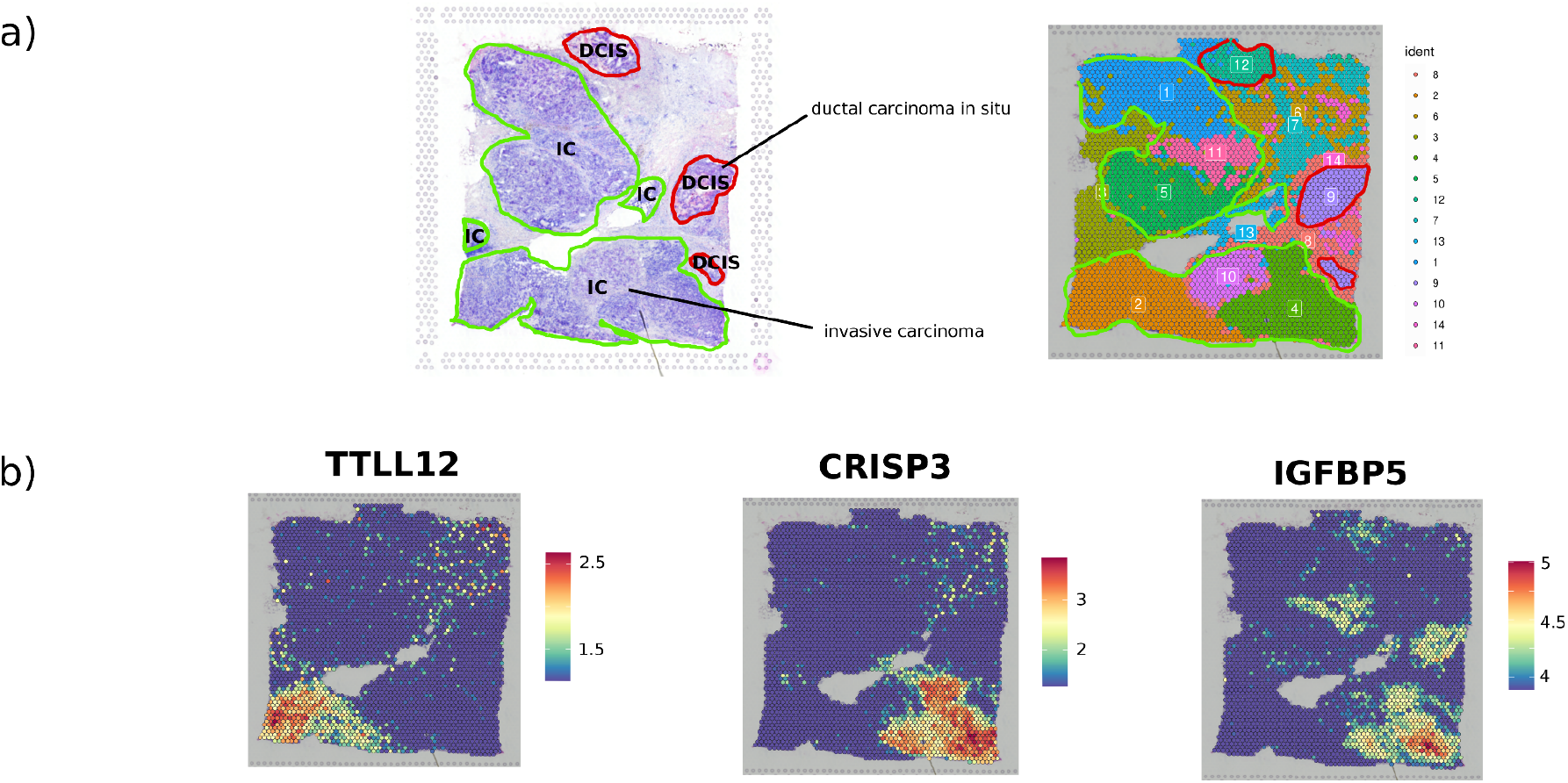
Cluster Biological coherence. (a) Manual pathologists annotation of human breast cancer 1 dataset provided by Lewis *et al*. [2] and clustering achieved with the best configuration of *Stardust\** with resolution 0.6. (b) Spatial plots showing the expression level of three of the top 100 genes with highest Moran Index for HBC1 Visium dataset.

We computed the Moran’s I also for breast cancer (HBC2), mouse kidney (MK), human heart (HH) and human lymph node (HLN) datasets (Supplementary file Figure S6). We observed that, as for HBC1 dataset, genes colocalize in well-defined cluster shapes attesting the quality of the results achieved by *Stardust\**.

*Stardust* and *Stardust\** were also tested and compared with state-of-the-art ST clustering methods, namely *stLearn, SpaGCN, Giotto* and *BayesSpace*, by analysing each individual 10x Genomics dataset. We evaluated *Stardust* stability scores with respect to the ones achieved with the other tools. We fixed the number of principal components to 10, which we found to be a good threshold for all the datasets analysed through the ‘*elbow*’ method proposed by rCASC [19]. The cluster resolution parameter was set to 0.6, 0.8 and 1 for each tool. For each resolution value, among the five configurations of *Stardust* obtained by varying the space weight parameter, we decided to represent the most stable ones, with space weight mainly equal to 0.25 and 0.5 for each dataset analysed. Since *BayesSpace* and *Giotto* require a priori knowledge on the number of clusters, we derived it from the results of *Stardust* using for each cluster resolution the *Stardust* configuration with the lowest coefficient of variation score. Moreover, we tested *SpaGCN* both including and excluding histology image information. Concerning HBC1, by looking at the coefficient of variation in Figure 5 (a) *Stardust* and *Stardust\** are the tools able to achieve the highest average stability score. Figure 5 (b) shows the stability results of the compared tools, using their best configurations according to Figure 5 (a), i.e. the ones with the lowest coefficient of variation value: resolution 0.6 and space weight 0.25 for *Stardust*, resolution 0.6 for *Stardust\**, resolution 0.6 and image True for *SpaGCN* and *stLearn*, resolution 0.8 for *BayesSpace* and resolution 0.6 for *Giotto. Stardust* and *Stardust\** reach the lowest coefficients followed by *stLearn* and *BayesSpace*. Results for *SpaGCN* with cluster resolution 1 are missing because computation was out of a predefined time (> 4h). Cluster results for a visual exploration together with the original tissue are shown in (Figure 5 (c)). According to the shifts of the stability scores (Figure 5 (d)), computed using threshold of 50%, *Stardust\** outperforms all other methods.

**Figure 5:**
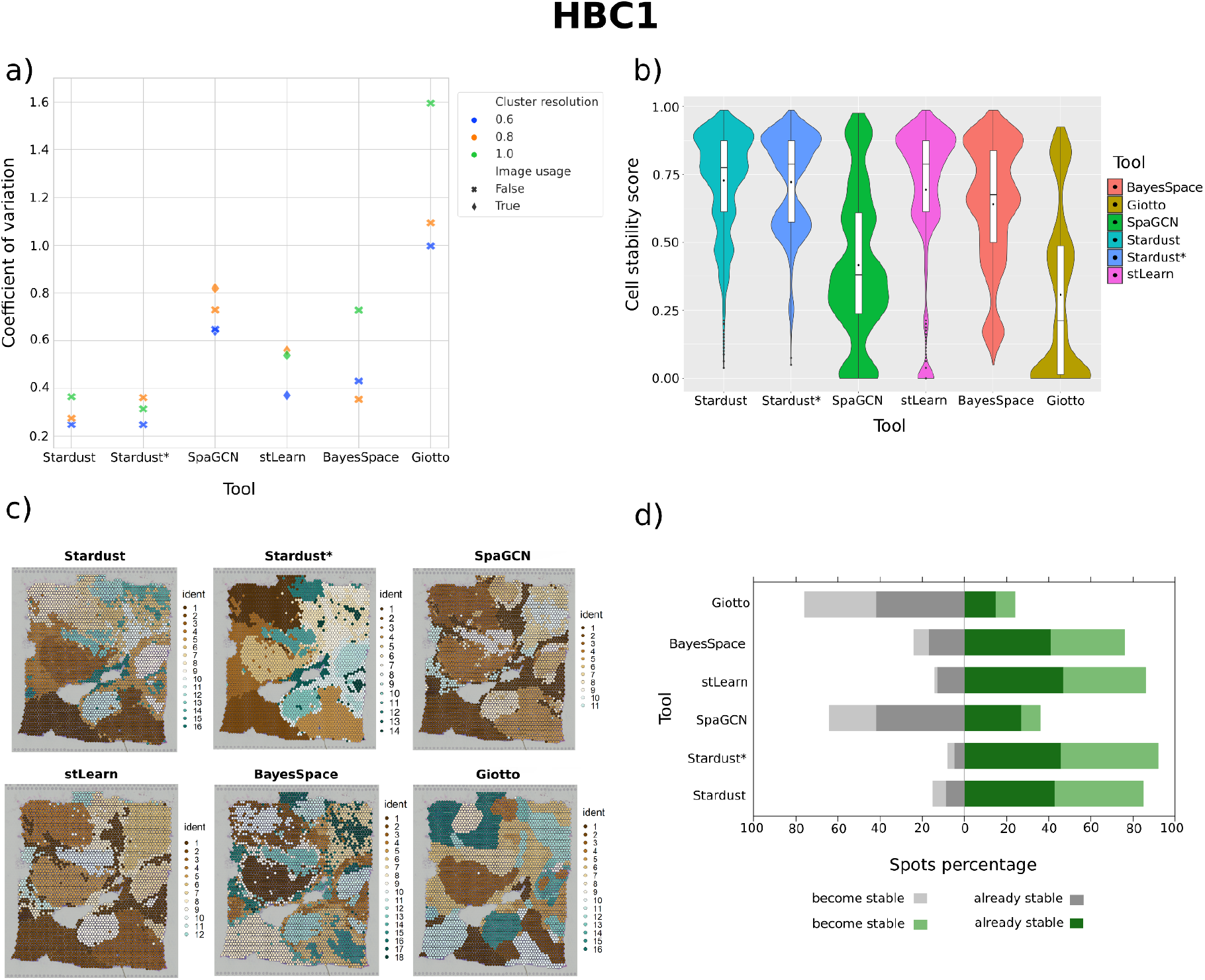
Comparison of *Stardust, Stardust\** and state of art tools on HBC1 dataset. (a) The coefficient of variation values derived from the stability score distribution of each tool configuration. The cluster resolution refers to the resolution parameter for the Louvain community detection algorithm, image usage tells whether the image is included in the clustering method. (b) The cell stability score distributions of the best performing configuration of each tool (i.e., the one with the lowest coefficient of variation). (c) The H&E (Hematoxylin & Eosin) stained tissue sample and a spatial plot for each best tool configuration with clusters of spots on the tissue section. (d) The stability scores shifts obtained comparing the best configuration of each tool with the base *Stardust no space* version, i.e., the one not considering space.

Analysing HBC2, HH, HLN, and MK (Supplementary file Figures S7-10), we observed that *Stardust* and *Stardust\** achieve the lowest values in terms of coefficient of variation (Figures S7-10 (a)) and the highest values in terms of average stability (Figures S7-10 (b)) overcoming all the other tools. In particular, we noticed that in some cases, only for HBC2, the coefficient of variations are comparable with *BayesSpace* (Figure S7 (a)). In MK (Figure S10 (a)), *Stardust* and *Stardust\** clearly show the lowest coefficient of variation value (Figure S10 (a)) and the highest stability scores, with an average value above 75% (Figure S10 (b)). Figures S7-10 (c) graphically show the formed clusters. In HBC2 and MK datasets, *Stardust* and *Stardust\** overcome all the other tools in terms of the highest percentage of spots that become stable (Figures S7 and S10 (d)). Our methods, as well as all the other tools, in tissues such as HH and HLN characterized by a more homogeneous architecture with respect to the morphological structure clearly visible in the other datasets (Figures S8 and S9 (d)), do not find a relevant number of spots showing a high degree of stability, except for *BayesSpace* that tends to find a slightly larger number of stable spots.

To show the scalability of *Stardust* we tested it on the five 10x datasets and on three datasets obtained from two different spatial sequencing technologies, namely Seq-scope [17] and Slide-seq [18]. Running time of *Stardust\** is equivalent to *Stardust*.

Supplementary Figure S11 shows that the relation between the size of the dataset and *Stardust* running time increases linearly. Although this linear relation, computational resources required for high-dimensional datasets, as those generated by Slide-seq technology, increment consistently. Quality of *Stardust\** clustering was also confirmed by analysing the biological consistency of the clusters in Seq-scope and Slide-seq datasets (Figures S12-S14 in Supplementary file) with the available cell annotations and by showing that features with highest Moran’s I colocalize inside clusters shapes.

## Conclusion

We developed *Stardust*, an open-source and easy to install R package for Spatial Transcriptomics data clustering which integrates transcriptional and spatial information through a complete auto-tuned approach. The package contains a method to manually explore the space influence on clustering (named as the package, *Stardust*) and one version fully automated called *Stardust\**. The tools performances were evaluated by analyzing the clustering stability through two stability measures: the cell stability score and the coefficient of variation. Moreover, we confirmed clustering biological coherence by comparing tissue architecture with cluster shapes and by computing the Moran’s I to identify the spatial autocorrelated features. Methods stability scores were compared with the ones achieved without using space to show how spatial information can significantly improve the clustering outcome. Results were also compared with those achieved by the state-of-the-art tools investigated, including *BayesSpace, SpaGCN, stLearn* and *Giotto*. Results of each dataset analysis assess that the proposed methods achieve more stable results with respect to clustering performed without considering spatial information and also that they are valid competitors, in terms of stability, to existing state-of-the-art clustering methods. Moreover, results demonstrated that the introduction of features from a histology image generally led to more unstable and misleading clustering results, particularly when the tissue section is quite uniform, and therefore, does not contain any particular structural information that could help clustering.

## Supporting information

Stardust_Supplementary_Section

## Availability of source code and requirements

- Project name: Stardust
- Project home page: https://github.com/InfOmics/stardust/, rCASC is available on https://github.com/InfOmics/rCASC and GenSA for Stardust is available on https://github.com/SimoneAvesani/Tuning_Stardust.
- Operating system(s): UNIX-like OS (MacOS or a Linux distribution)
- Programming language: R
- Other requirements: Docker
- License: MIT license
- Any restrictions to use by non-academics: None

## Availability of supporting data

10x datasets are available at https://github.com/InfOmics/stardust/. After the registration on the 10x Genomics website, each individual dataset can be downloaded from:

- Human breast cancer (HBC1): https://www.10xgenomics.com/resources/datasets/human-breast-cancer-block-a-section-1-1-standard-1-1-0
- Human breast cancer (HBC2): https://www.10xgenomics.com/resources/datasets/human-breast-cancer-block-a-section-2-1-standard-1-1-0
- Human heart (HH): https://www.10xgenomics.com/resources/datasets/human-heart-1-standard-1-1-0
- Human lymph node (HLN): https://www.10xgenomics.com/resources/datasets/human-lymph-node-1-standard-1-1-0
- Mouse kidney (MK): https://www.10xgenomics.com/resources/datasets/mouse-kidney-section-coronal-1-standard-1-1-0

Seq-scope datasets are available at the Deep Blue Data platform: https://deepblue.lib.umich.edu/data/concern/data_sets/9c67wn05f?locale=en.

- Colon: https://deepblue.lib.umich.edu/data/downloads/rb68xc160
- Liver *TD*: https://deepblue.lib.umich.edu/data/downloads/7w62f844w

Slide-seq cerebellum dataset is available, after registration, at the Broad Institute Single Cell Portal: https://singlecell.broadinstitute.org/single_cell/study/SCP948.

## Abbreviations

CSS: Cell Stability Score
DCIS: Ductal Carcinoma In Situ
DLPFC: Dorsolateral Prefrontal Cortex
GenSA: Generalized Simulated Annealing
HBC1: Human Breast Cancer 1
HBC2: Human Breast Cancer
HH: Human Heart
HLN: Human Lymph Node
HMRF: Hidden Markov Random Field
H&E: Hematoxylin & Eosin
IC: Invasive Carcinoma
KNN: K-Nearest Neighbor
MCMC: Markov chain Monte Carlo
MK: Mouse Kidney
MRF: Markov Random Field
PC: Principal Component
PCA: Principal Component Analysis
rCASC: reproducible Classification Analysis of Single Cell Sequencing Data
RDS: R Data Serialized
scRNA-seq: Single-cell RNA sequencing
ST: Spatial Transcriptomics

## Author contributions

Conceptualization: RG and GM; Methodology: RG, SA, EV, LA, GM, VB, MB, RC; Supervision: RG, RC; Writing: RG, SA, EV; Review & editing: all; Code Writing: SA, EV, LA, GM; Test: SA, EV; Validation: RG, SA, EV, RC. None of the authors have any competing interests in the manuscript.

## Acknowledgments

We thank prof. Giorgio Cattoretti for the useful comments and discussions on the paper.

## Supplementary Material

- Supplementary material file name : *Stardust_Supplementary_Section*
- Extension: .*pdf*
- Description: *Stardust tests*

