## Supplementary material for "Stardust: improving spatial transcriptomics data analysis through space aware modularity optimization based clustering": Stardust_Supplementary_Section

* equal contributor
### equal contributor

#### Analysis of the influence of space information in *Stardust* clustering

Figure S1 showcases the true power of exploiting spatial information to achieve better stability scores when clustering Spatial Transcriptomic data. When space is not considered like in clusters plot with title space weight 0.0 almost none of the spots can achieve better stability scores than the threshold of 0.5. In the best *Stardust* configuration (space weight 1), the majority of them become stable.

An example, in particular, of increased spatial coherence in this dataset is represented by the area A circled in red in the tissue image in the Figure S1 (a) related to configuration space weight 0.0. Clusters 3 (spots depicted in white) and 6 (spots depicted in light brown) are located mainly in that area and are spatially close. They achieve poor stability scores as shown on the circled area C of the corresponding images in Figure S1 (b), due to high variability of cluster identity assignment to spots. *Stardust* collapses area A in one cluster (area B in Figure S1 (a)) that is spatially well-defined and achieves high stability scores (area D in Figure S1 (b)).

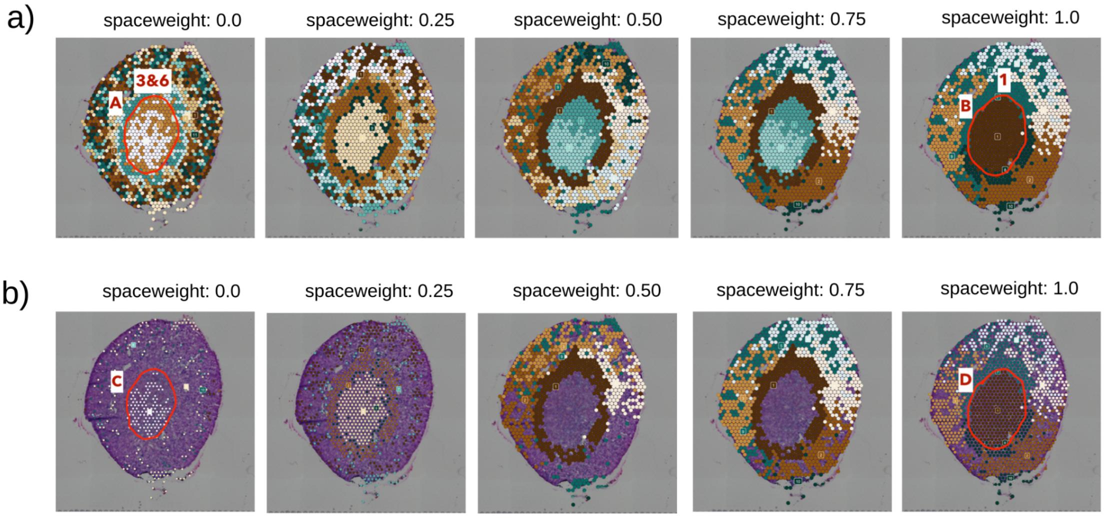


**Figure S1: Spatial clusters plots in mouse kidney (MK) dataset. In (a) the results of 5 *Stardust* configurations with increasing space weight are shown. The same results, where only spots that have stability score >= 0.5 are visualized, are shown in (b). Each color corresponds to one of the 11, 9, 10, 10 and 10 cluster identities obtained in each configuration (in order of appearance), respectively.**

In Figure S2 we show that the effect of introducing spatial information in the clustering task for the human lymph node dataset, other than increasing the stability scores, is the mitigation of cases in which a subset of spots is not recognized as a unique cluster and is separated into two or more unstable clusters. An example of increased spatial coherence in this dataset is represented by the area A circled in red in the tissue image in Figure S2 (a) related to configuration space weight 0.0, i.e. when no space is used. Clusters 7 (spots depicted in white) and 9 (spots depicted in light cyan) are located mainly in that area and are spatially close. They achieve poor stability scores as shown on the circled area C of the corresponding images in Figure S2 (b), due to the high variability of cluster identity assignment to spots. The best performing *Stardust* configuration collapses area A in one cluster (area B in Figure S2 (a) configuration space weight 0.50 ) that is spatially well-defined and achieves high stability scores (area D in Figure S2 (b) on the same configuration).

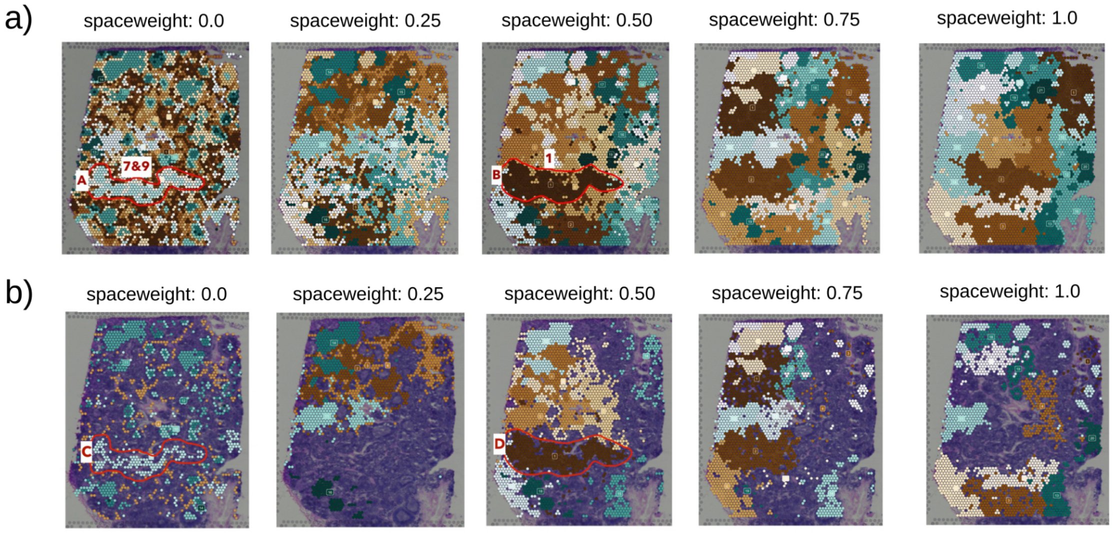


**Figure S2: Spatial clusters plots in human lymph node (HLN) dataset. In (a), the results of 5 *Stardust* configurations with increasing space weight are shown. The same results, where only spots that have stability score >= 0.5 are visualized, are shown in (b). Each color corresponds to one of the 14, 16, 19, 21 and 21 cluster identities obtained in each configuration (in order of appearance), respectively.**

Visualizations in Figure S3 demonstrate that *Stardust* does not create a structure where it's not present. In fact, human heart (HH) tissue presents a similar architecture across the whole sample and the usage of space doesn't evidentiate a relevant number of stable subgroups of spots (clusters). The more evident results that *Stardust* is able to obtain are represented by the areas A and B circled in red in the tissue image in Figures S3 (a) and (b) related to configuration space weight 0.50. Although clusters 9 and 1 achieve good stability, they cover only a small portion of the tissue confirming that *Stardust* needs transcriptional variability across the tissue to identify structures in the tissue architecture.

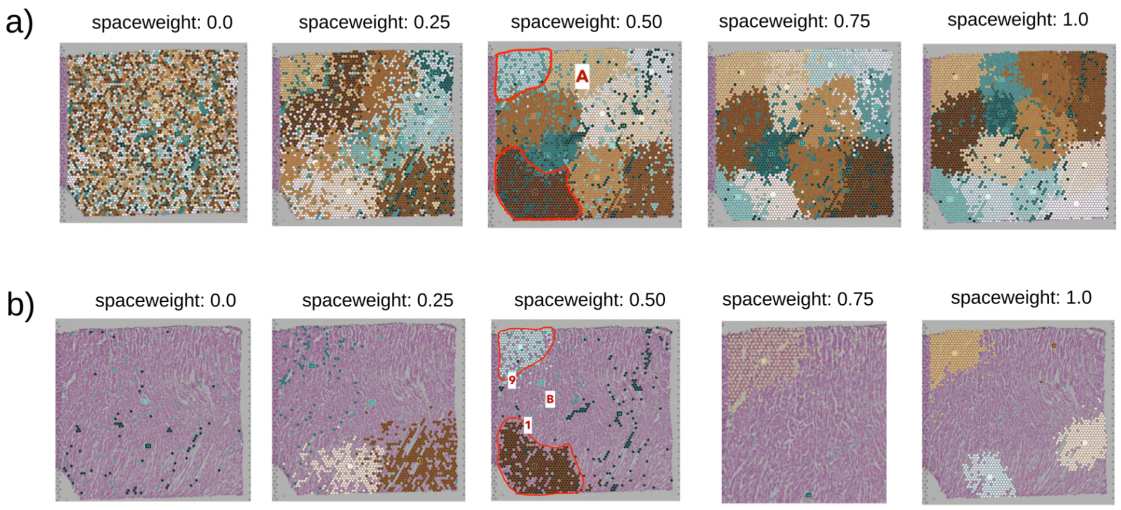


**Figure S3: Spatial clusters plots in human heart (HH) dataset. In (a), the results of 5 *Stardust* configurations with increasing space weight are shown. The same results, where only spots that have stability score >= 0.5 are visualized, are shown in (b). Each color corresponds to one of the 13, 15, 14, and 15 cluster identities obtained in each configuration (in order of appearance), respectively.**

In Figure S4, related to the human breast cancer (HBC1) dataset, spatial information helps to mitigate cases in which distant spots are clustered together due to transcriptional similarity, but they do not belong to the same region in the tissue architecture. An example is represented by area A circled in red in the first cluster plot in Figure S4 (a). Area A contains the clusters 6 (spots in beige). When clusters 6 are considered as a whole as when space is not considered (i.e. configuration space weight 0.0), it achieves poor stability scores as shown on the circled area C (Figure S4 (b)). The best performing *Stardust* configuration (space weight 0.75) divides area A into two clusters (16, 14 in area B) that achieve high stability scores (area D). This is a notable result of *Stardust*, infact in [2] the authors report a manual annotation of the same tissue where the two clusters are identified as DCIS (ductal carcinoma in situ) compared to other clusters that have been defined as IC (invasive carcinoma). If the no use of space allows identifying the clusters 6 as a single cluster, these clusters are unstable (Figure S4 (b)) and therefore in unknown situations they would not be suggested to the pathologist to be noticed. Indeed, *Stardust* in its best configuration identifies them as very stable, albeit separate, clusters. The separation is foreseen as they are distant.

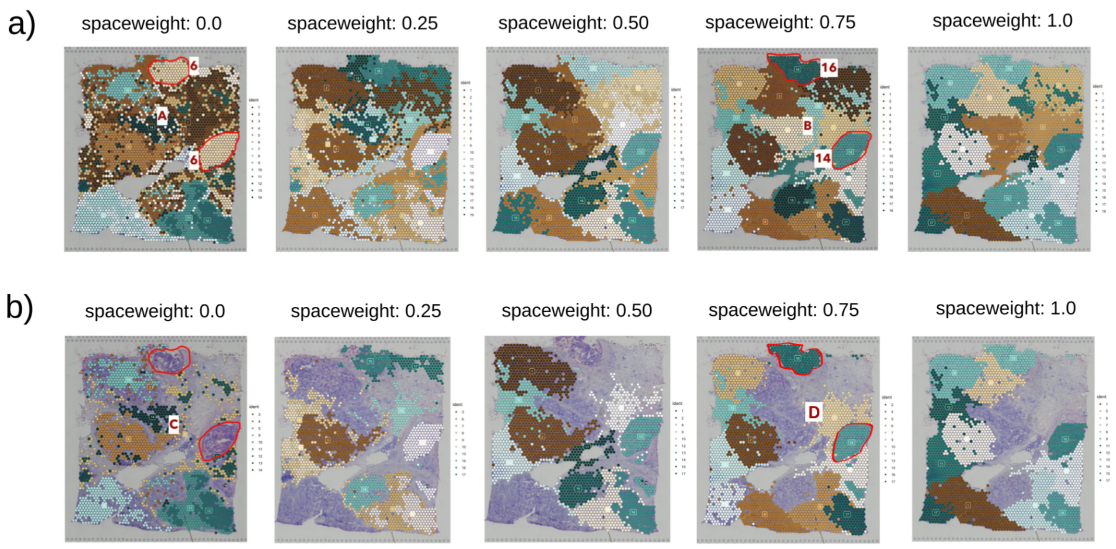

**Figure S4: Spatial clusters plots in human breast cancer (HBC1) dataset. In (a), the results of 5 *Stardust* configurations with increasing space weight are shown. The same results, where only spots that have stability score >= 0.5 are visualized, are shown in (b). Each color corresponds to one of the 14, 19, 17, 18 and 18 cluster identities obtained in each configuration (in order of appearance), respectively.**

In Figure S5, related to the human breast cancer (HBC2) dataset, spatial information increases the overall stability of two spatially neighbouring clusters. In this case, the strategy adopted by *Stardust* to increase stability scores is not to merge them as in HLN dataset but to reassign the cluster identity of each spot keeping two spatially neighbouring clusters. In the red circled area A in Figure S5 (a) when no space is used (i.e. configuration space weight 0.0) is depicted the original cluster arrangement that performs poorly (area C in Figure S5 (b)). In red circled area B (Figure S5 (a) configuration space weight 0.50) is shown the *Stardust* rearrangement with increased stability scores (area D in Figure S5 (b)).

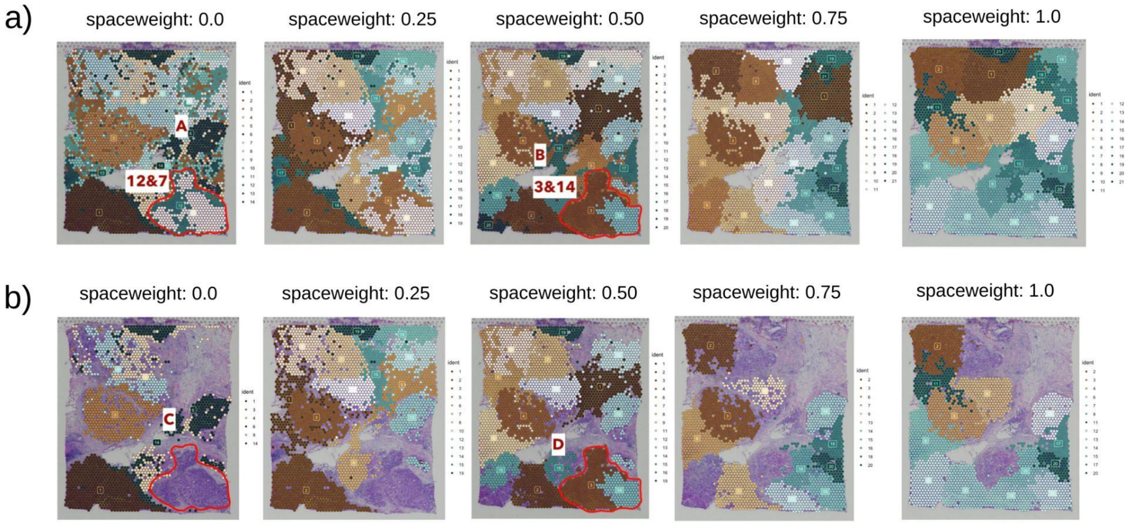


**Figure S5: Spatial clusters plots in human breast cancer (HBC2) dataset. In (a), the results of 5 *Stardust* configurations with increasing space weight are shown. The same results, where only spots that have stability score >= 0.5 are visualized, are shown in (b). Each colour corresponds to one of the 14, 19, 20, 21 and 21 cluster identities obtained in each configuration (in order of appearance), respectively.**

We investigated the biological coherence of the obtained *Stardust** clusters by computing the Moran's I. Moran's I uses both feature locations and feature values simultaneously to define the genes' spatial autocorrelation. Figure S6 reports the Moran's I on breast cancer (HBC2), mouse kidney (MK), human heart (HH) and human lymph node (HLN) datasets.

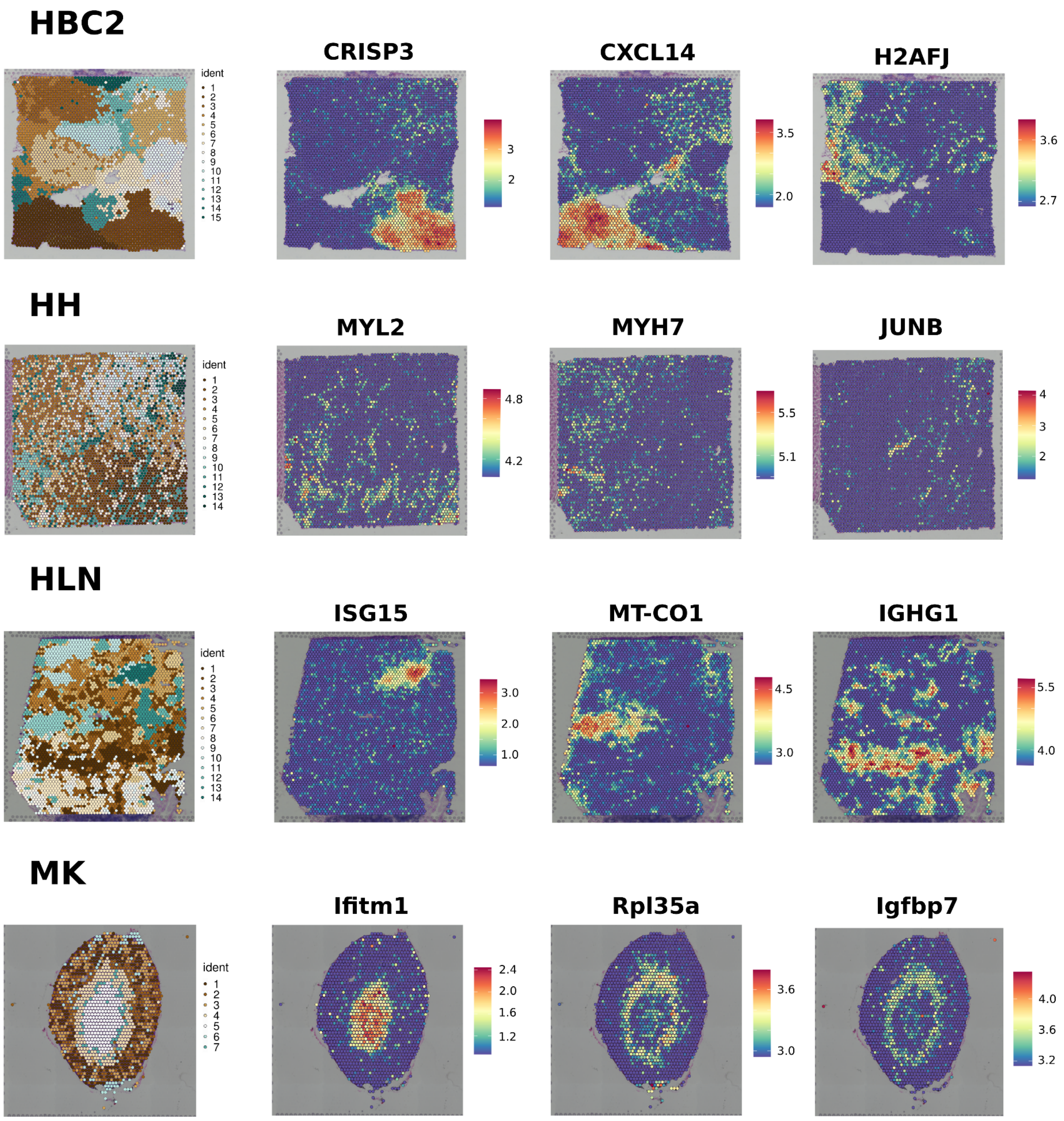


**Figure S6: Spatial plots showing the expression level of three of the top 100 genes with highest Moran's index for HBC2, HH, HLN and MK Visium datasets and the clusters obtained with *Stardust****.**

### Comparing *Stardust* and *Stardust** with existing tools

Figure S7 reports details on the analysis of HBC2. The best configurations for achieving the lowest coefficient of variation value are (Figure S7 (a)): resolution 0.6 and space weight 0.5 for *Stardust*, resolution 0.6 for *Stardust*,* resolution 0.6 and image False for *SpaGCN*, resolution 1 for *stLearn* and *BayesSpace* and resolution 0.6 for *Giotto*. Results for *SpaGCN* with cluster resolution 0.8 without image and 1 are missing because computation was out of a predefined time (> 4h). *Stardust** outperforms all the other tools, including *BayesSpace,* in terms of average cell stability score (Figure S7 (b)). Cluster results for a visual exploration are shown in (Figure S7 (c)). The analysis of shift scores (Figure S7 (d)) confirms the clustering stability of *Stardust** followed by *BayesSpace.*

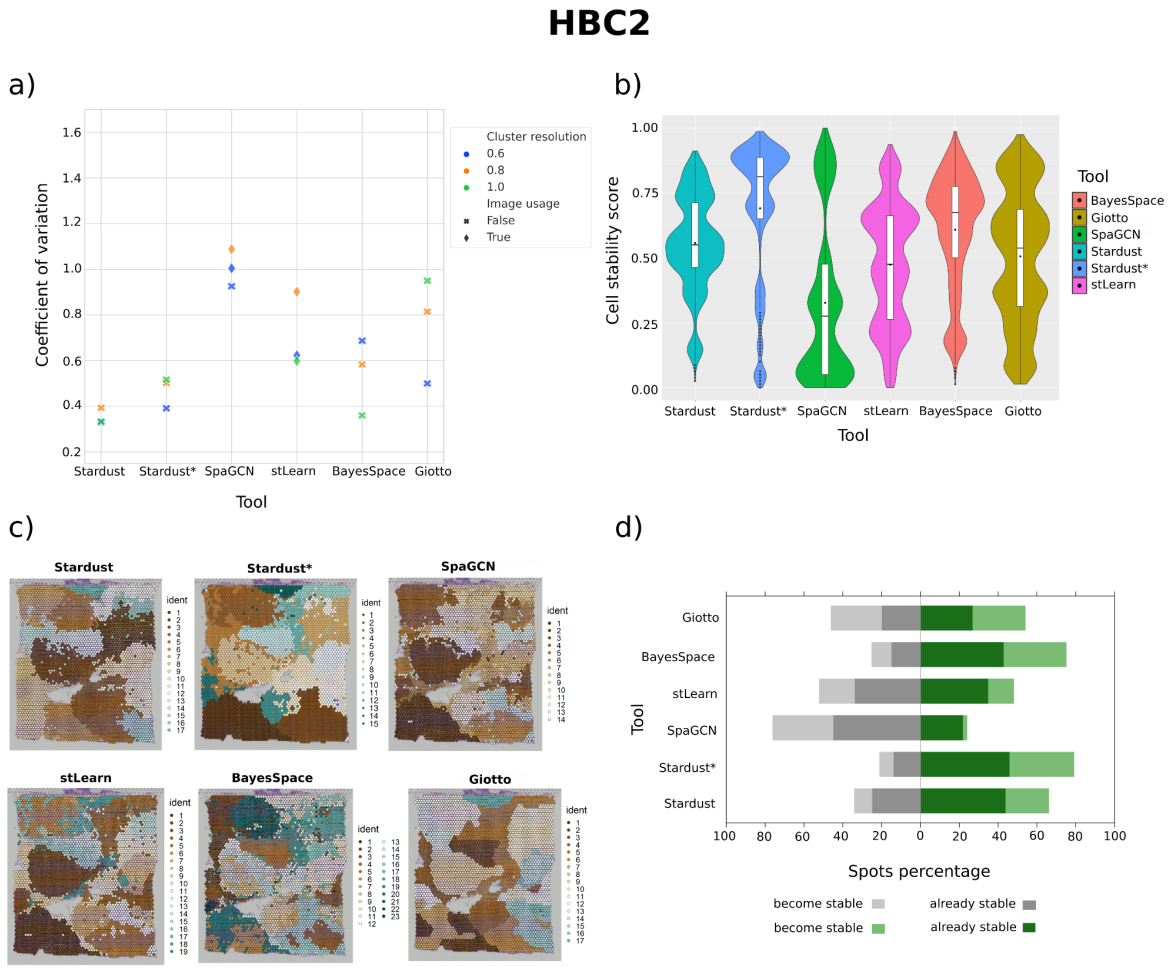


**Figure S7: Comparison of *Stardust*, *Stardust** and state of art tools on HBC2 dataset: (a) The coefficient of variation values derived from the stability score distribution of each tool configuration. The cluster resolution refers to the resolution parameter for the Louvain community detection algorithm, image usage tells whether the image is included in the clustering method. (b) The cell stability score distributions of the best performing configuration of each tool (i.e., the one with the lowest coefficient of variation). (c) The H&E (Hematoxylin & Eosin) stained tissue sample and a spatial plot for each best tool configuration with clusters of spots on the tissue section. (d) The stability scores shifts obtained comparing the best configuration of each tool with the base *Stardust no space* version, i.e., the one not considering space.**

In HH analysis (Figure S8), the best configurations achieving the lowest coefficient of variation value are (Figure S8 (a)): resolution 0.8 and space weight 0.5 for *Stardust*, resolution 1 for *Stardust**, resolution 0.6 and image False for *SpaGCN*, resolution 0.6 for *stLearn*, resolution 0.8 for *BayesSpace* and resolution 1 for *Giotto*. Results for *SpaGCN* with resolution 1 with the use of image are not reported because it is an out-of-scale value. Looking at the stability score distributions and at the score shifts, *Stardust** is the tool with the highest average stability (Figure S8 (b)). Cluster results for a visual exploration are shown in (Figure S8 (c)). *BayesSpace* shows higher percentage of spots becoming stable (Figure S8 (d)), followed by *Stardust**. However, HH tissue demonstrates that *Stardust* and Stardust* are able to avoid finding deceptive structures when a well-defined histological pattern is not present.

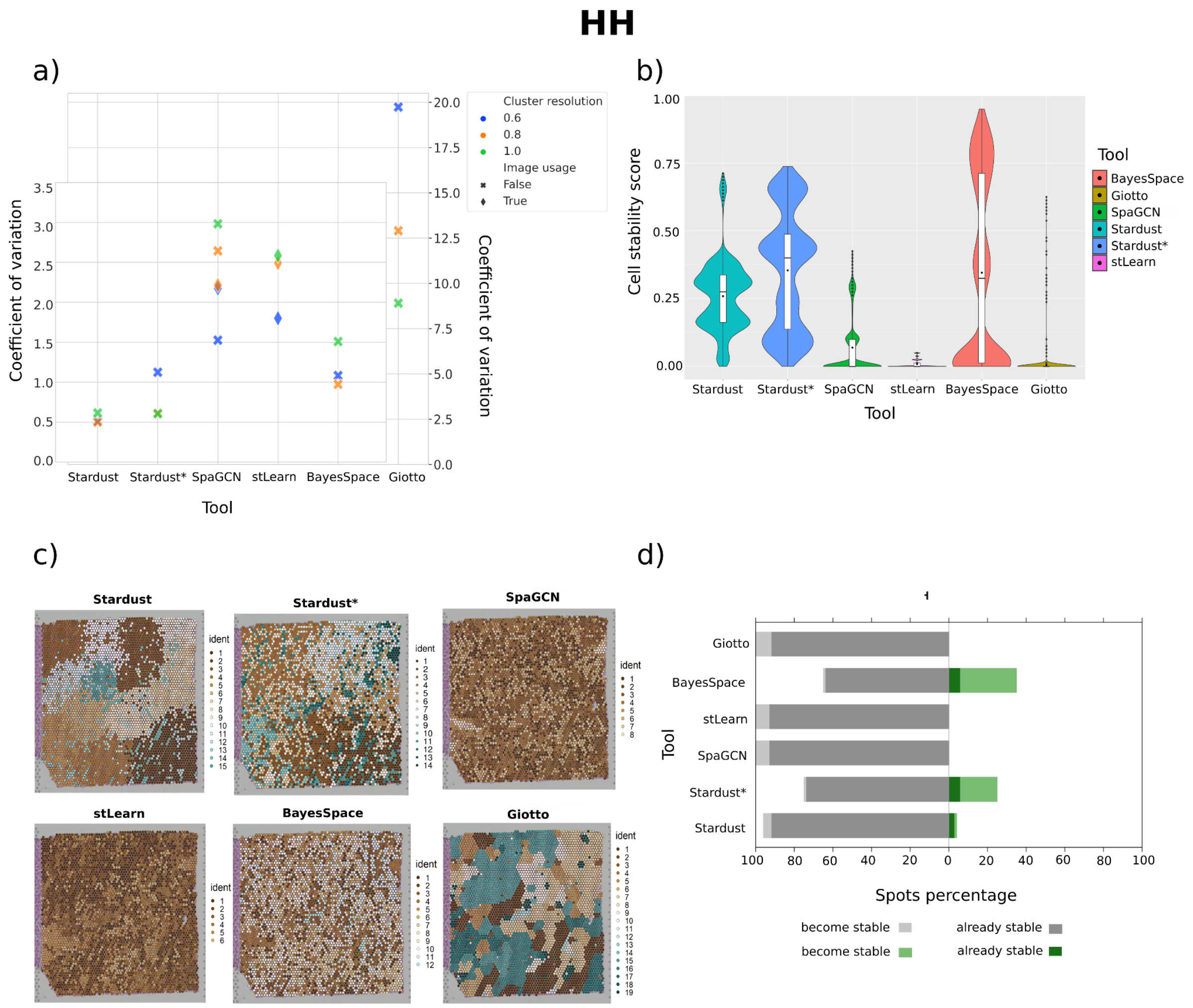


**Figure S8: Comparison of *Stardust* and *Stardust** and state of art tools on HH dataset. (a) The coefficient of variation values derived from the stability score distribution of each tool configuration. The cluster resolution refers to the resolution parameter for the Louvain community detection algorithm, image usage tells whether the image is included in the clustering method. (b) The cell stability score distributions of the best performing configuration of each tool (i.e., the one with the lowest coefficient of variation). (c) The H&E (Hematoxylin & Eosin) stained tissue sample and a spatial plot for each best tool configuration with clusters of spots on the tissue section. (d) The stability scores shifts obtained comparing the best configuration of each tool with the base *Stardust no space* version, i.e., the one not considering space.**

Figure S9 reports details of HLN analysis. The best configurations achieving the lowest coefficient of variation value are (Figure S9 (a)): resolution 0.8 and space weight 0.25 for *Stardust*, resolution 0.8 for *Stardust**, resolution 0.6 and image False for *SpaGCN*, resolution 0.8 for *stLearn*, resolution 0.6 for *BayesSpace* and *Giotto*. Results for *SpaGCN* with cluster resolution 1 are not reported because computation was out of a predefined time (> 4h). *Stardust* exceeds the stability scores of the other tools (Figure S9 (b)). Cluster results for a visual exploration are shown in (Figure S9 (c)). *BayesSpace* achieves good results, followed by our methods while the other tools show very low stability scores, especially *Giotto*, in which there are no spots that become stable (Figure S9 (d)).

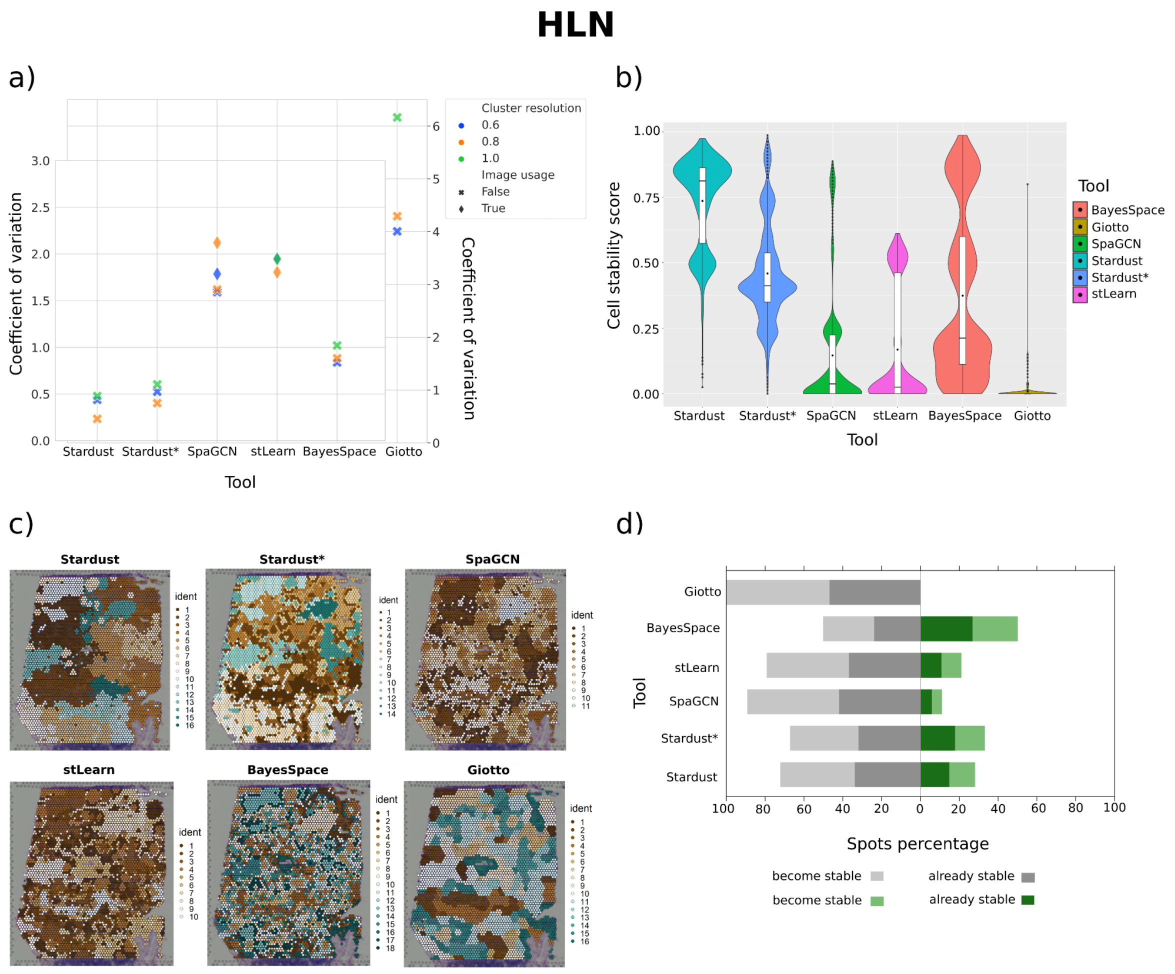


**Figure S9: Comparison of *Stardust*, and *Stardust** and state of art tools on HLN dataset. (a) The coefficient of variation values derived from the stability score distribution of each tool configuration. The cluster resolution refers to the resolution parameter for the Louvain community detection algorithm, image usage tells whether the image is included in the clustering method. (b) The cell stability score distributions of the best performing configuration of each tool (i.e., the one with the lowest coefficient of variation). (c) The H&E (Hematoxylin & Eosin) stained tissue sample and a spatial plot for each best tool configuration with clusters of spots on the tissue section. (d) The stability scores shifts obtained comparing the best configuration of each tool with the base *Stardust no space* version, i.e., the one not considering space.**

In MK analysis (Figure S10), the best configurations achieving the lowest coefficient of variation value are (Figure S10 (a)): resolution 0.6 and space weight 0.25 for *Stardust*, resolution 0.6 for *Stardust**, resolution 0.6 and image True for *SpaGCN*, resolution 0.8 for *stLearn*, resolution 1 for *BayesSpace* and resolution 0.6 for *Giotto*. Spatial clustering appears similar in almost all methods (Figure S10 (c)), except for *Giotto* for which the clustering stability is indeed significantly lower as compared to the other tools (Figure S10 (d)). The proposed methods overcome all other tools in terms of the highest percentage of spots that become stable (Figure S10 (d)).

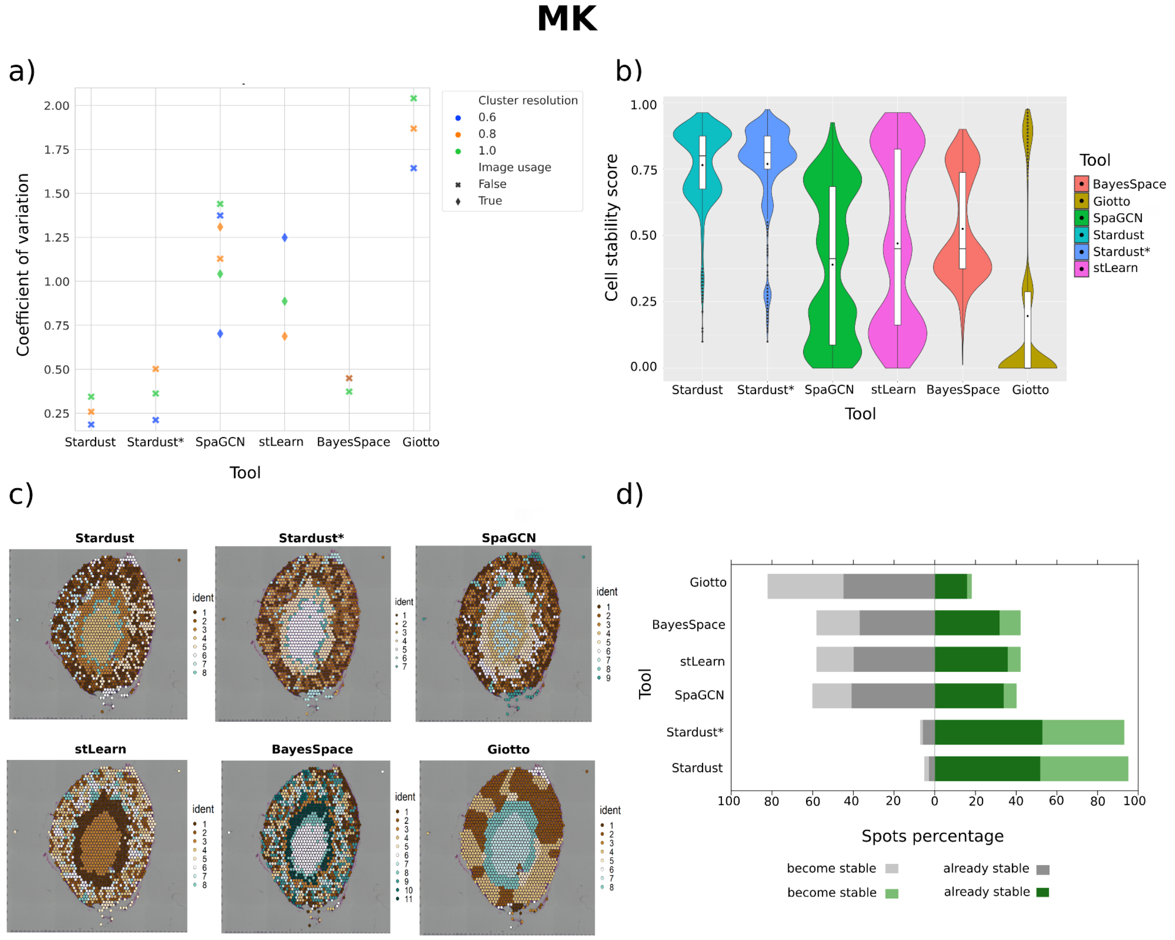


**Figure S10: Comparison of *Stardust*, *Stardust**, and state of art tools on MK dataset. (a) The coefficient of variation values derived from the stability score distribution of each tool configuration. The cluster resolution refers to the resolution parameter for the Louvain community detection algorithm, image usage tells whether the image is included in the clustering method. (b) The cell stability score distributions of the best performing configuration of each tool (i.e., the one with the lowest coefficient of variation). (c) The H&E (Hematoxylin & Eosin) stained tissue sample and a spatial plot for each best tool configuration with clusters of spots on the tissue section. (d) The stability scores shifts obtained comparing the best configuration of each tool with the base *Stardust no space* version, i.e., the one not considering space.**

### *Stardust* and *Stardust** scalability and biological coherence

We explored the time scalability of proposed methods (Figure S11) on five 10x, two Seq-scope and one Slide-seq datasets. Data were selected to have a different number of spots/cells. Running time of *Stardust* and *Stardust** are equivalent. Methods scale linearly with the dimension of the datasets.

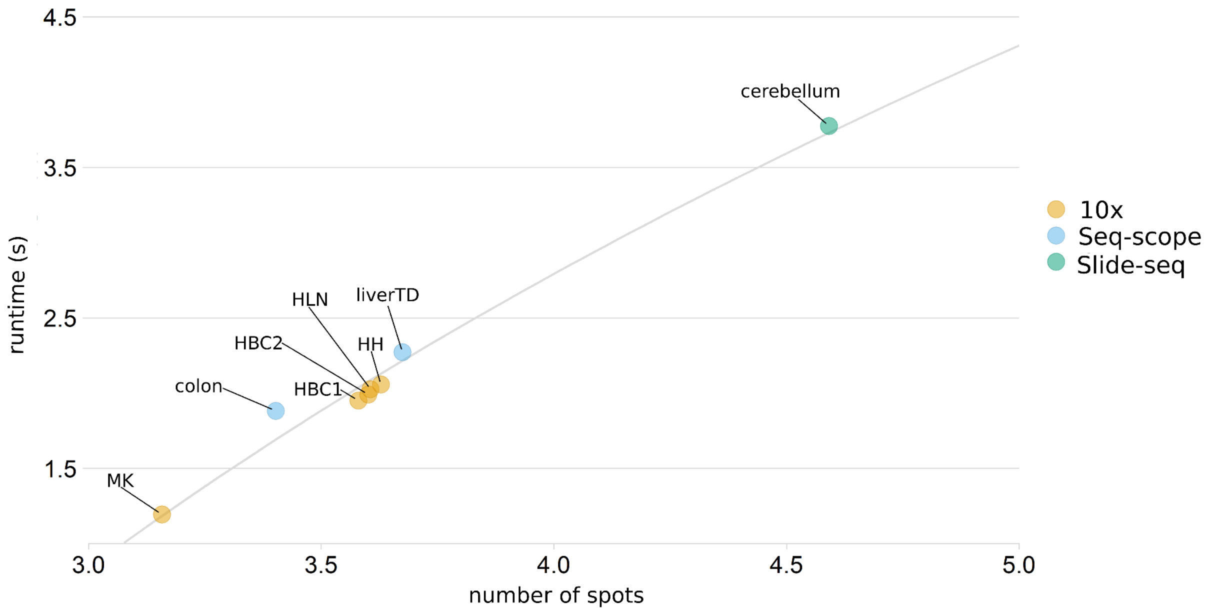


**Figure S11: Time scalability of *Stardust* on five 10x datasets, two Seq-scope datasets and one Slide-seq dataset. Axes values are in the log10 scale.**

We completed the verification of the biological consistency of *Stardust* and *Stardust** also on the clustering results obtained from the analysis of two Seq-scope datasets and one Slide-seq dataset. In order to do that we compared the achieved clusters with the provided cell type annotations and we computed the Moran's index to explore gene spatial autocorrelation (Figures S12-14). Notice that annotations for Slide-seq are available on a subset of the original dataset.

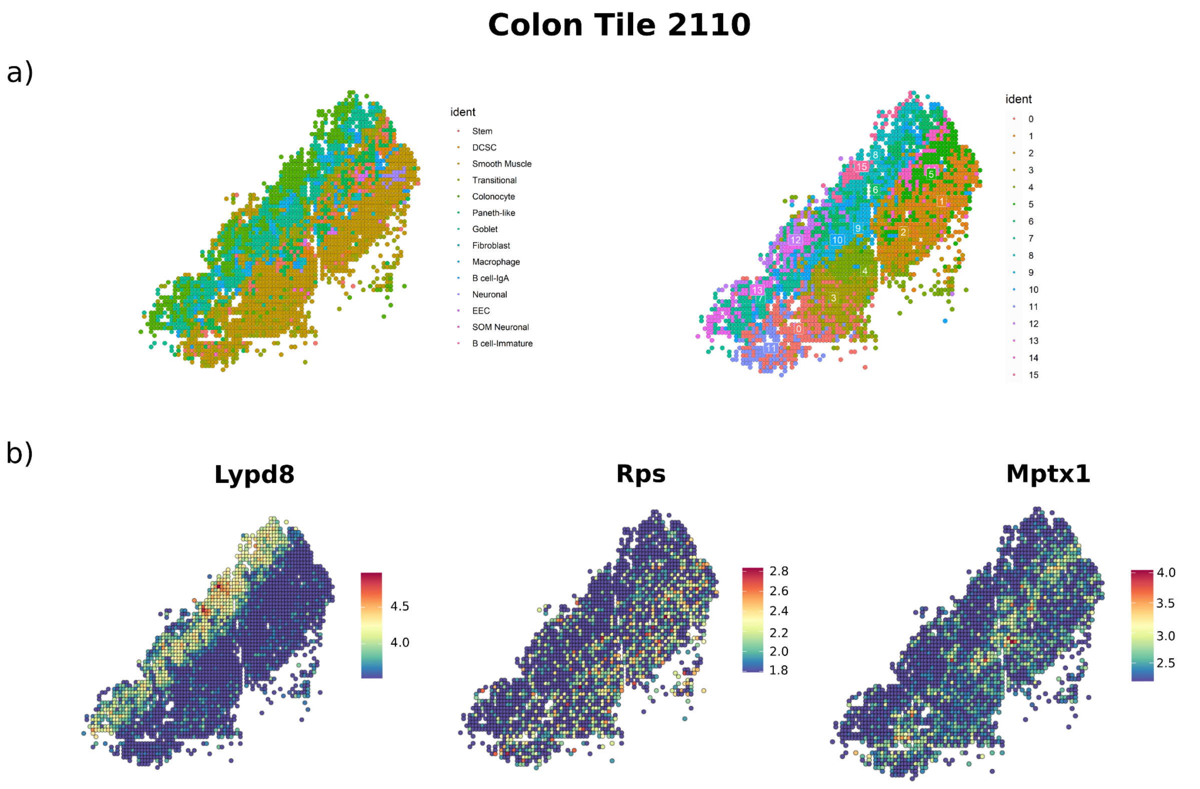


**Figure S12: Cluster Biological coherence.** (a) Cell type annotation of Seq-scope Colon Tile 2110 dataset and clustering achieved using *Stardust** with cluster resolution 1.5 as in [21]. (b) Spatial plots showing the expression level of three of the top 100 genes with highest Moran's I for Seq-scope Colon Tile 2110 dataset.

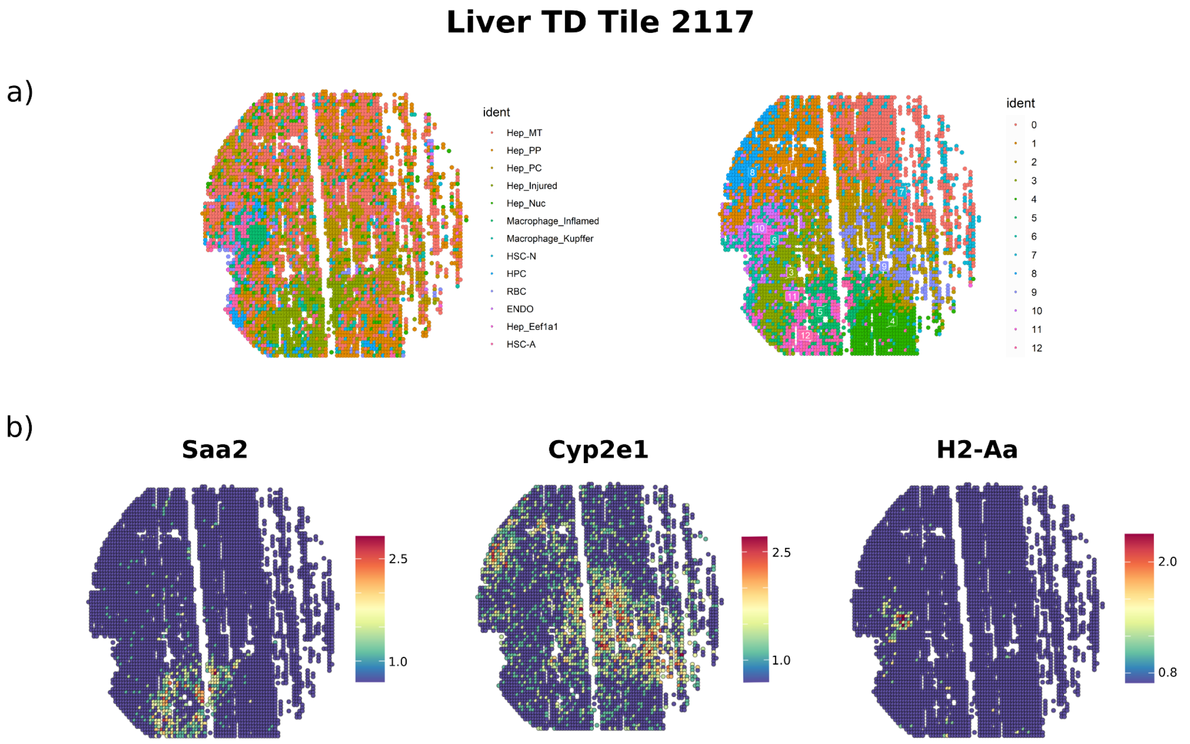


**Figure S13: Cluster Biological coherence. (a) Cell type annotation of Seq-scope Liver *TD* Tile 2117 dataset and clustering achieved using *Stardust** with cluster resolution 1 as in [18]. (b) Spatial plots showing the expression level of three of the top 100 genes with highest Moran's I for Seq-scope Liver *TD* Tile 2117 dataset.**

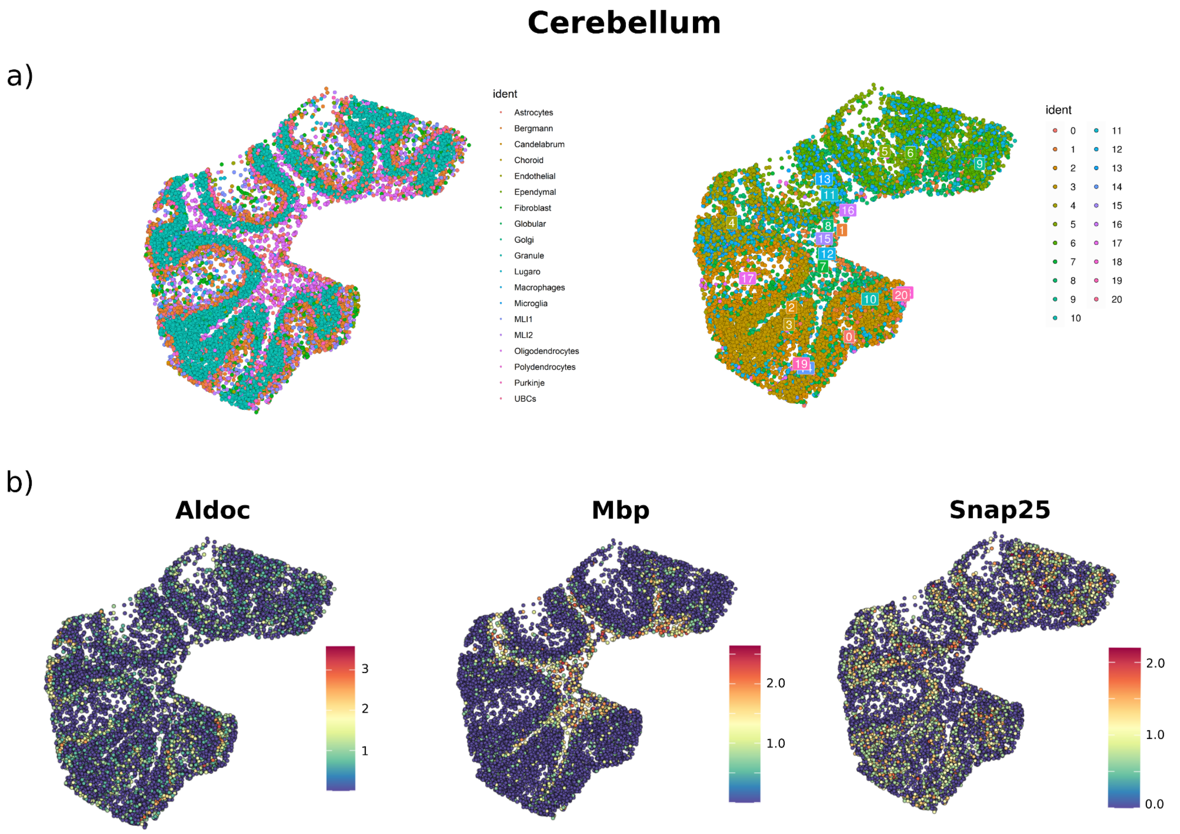


**Figure S14: Cluster Biological coherence. (a) Cell type annotation of Slide-seq cerebellum dataset and clustering achieved using *Stardust** with cluster resolution 0.6. (b) Spatial plots showing the expression level of three of the top 100 genes with highest Moran's I for Slide-seq cerebellum dataset.**
